# Phylogenetic comparative analyses of the determinants of anti-predator distraction behavior in shorebirds

**DOI:** 10.1101/2020.06.07.138537

**Authors:** Miki Saijo, Nobuyuki Kutsukake

## Abstract

Predation risk exerts a strong selective pressure on anti-predator behavior, resulting in behaviors to achieve defense of offspring and the individual. In shorebirds, some species perform distraction behavior that is attracting the attention of a predator. This behavior evolved, and were lost multiple times, independently and the behavioral repertoire varies among species. Although defense of offspring is critical for parents, the determinants of inter-specific variation in the distraction behavior remain unstudied. We surveyed the literature and conducted phylogenetic comparative analyses (n = 169 species) to test predictions regarding nest site, body mass, and coloniality. We found that small species were more likely to perform distraction behavior than large species. Solitary species were more likely to perform distraction behavior than colonial nesting species. Previous studies suggested that colonial nesting and large species commonly perform aggressive anti-predator behavior, implying that distraction behavior is an alternative anti-predator strategy to aggressive ones.

## Introduction

Predation risk exerts a strong selection pressure on animal behaviors, producing anti-predator behavior among others (Lack 1954; Caro 2005). Offspring predation is the most important reason for nest failure in birds (Clark & Wilson, 1981). Therefore, parents have evolved various anti-predator behaviors to counter offspring predation, thus increasing their fitness (Royle et al. 2012).

Shorebirds (Charadriiformes) are an ideal taxon for assessing the determinants of the anti-predator behavioral repertoire. Most shorebird species nest on open ground, exposing the chicks and eggs to strong predation pressure (Gochfeld, 1984; Kubelka et al., 2018). Shorebirds engage in two broad types of anti-predator behavior: attack and distraction (Gochfeld, 1984). Attack behavior includes any type of attack on predators, such as mobbing and scolding. Distraction behavior aims to attract the attention of a predator via injury-feigning, false breeding, and “rodent run” behaviors, for example (Caro, 2005; Gochfeld, 1984). When a predator was deceived by the display and chasing a parental bird, the bird moves away from their offspring (Ristau, 1991). As a result, predators lost sight of the nest or chick. When the predator was lured and far away from the nest by the display, the parental bird ends its display and runs away (Gómez-Serrano, 2018). This behavior seems to have evolved multiple times independently under similar ecological and social conditions (Humphreys & Ruxton, 2020). However, only a few studies have examined the determinants of inter-species differences in the anti-predator behavioral repertoire (Larsen et al. 1996; da Cunha et al. 2017). Larsen et al. (1996) tested factors of aggressive nest protection behavior in waders. This paper revealed that body mass and number of parents present on the nest territory explain the variation of aggressive anti-predator behavior. But they did not use phylogenetic comparative analysis. da Cunha et al. (2017) showed that small species were more likely to perform aggressive mobbing than large species. They included anti-predator behavior during both the breeding and non-breeding seasons.

In contrast to those studies analyzing aggressive anti-predator behavior, the determinants of distraction behavior remained unstudied. Armstrong (1954) listed six factors that could explain the performance of distraction behavior, which could be summarized that distraction display is likely to evolve in species that may be preyed upon by terrestrial predators during the daytime (Humphreys & Ruxton, 2020). However, a formal test of those ideas using a modern phylogenetic comparative analysis has not been done. We assessed three possible determinants of distraction behavior: nesting site, body mass, and coloniality. First, we predicted that species that nest in trees and on cliffs are less likely to perform anti-predator behavior, because their nests are less likely to be predated due to the difficulty of access compared with ground-nesting species (Coulson & Thomas, 1985). Differences in other traits according to nesting site were suggested in a detailed comparative study of two closely related sympatric species (Cullen, 1957). Cliff-nesting kittiwakes have weak anti-predator traits, such as rare production of alarm calls, unconcealed body color of chicks, and low response to an attack by a predator. By contrast, the parents of the ground-nesting black-headed gull (*Larus ridibundus*) attack predators, and their chicks escaped when attacked by a predator. We predicted that this behavioral difference between the two species is commonly seen in other taxa. Second, we predicted that body size would affect distraction behavior. Because of the physical advantages of large species, it has been supported such species are more likely to perform attack behavior than smaller species (Larsen et al. 1996), whereas smaller species are more likely to perform distraction behavior. Third, we predicted that solitary nesting species are more likely to perform distraction behavior because they cannot attack predators effectively compared with colonial species. Previous studies showed that colonial species are likely to perform attack behavior because they can attack a predator effectively as a group (Hoogland & Sherman, 1976; Robinson, 1985) and because they can minimize the per capita risk of predation via the dilution effect (Hamilton, 1971; Hogan, Hildenbrandt, Scott-Samuel, Cuthill, & Hemelrijk, 2017).

## Methods

### Data collection

We searched the literature for studies on anti-predator behavior during the breeding season using Google Scholar, with the key words of ‘shorebirds’, ‘plover’, ‘sandpiper’, ‘wader’, ‘gull’, ‘anti-predator’, ‘behavior’, and ‘nesting’, and the scientific or species names of Charadriiformes. We also added the key words ‘distraction’, ‘injury-feigning’, ‘false-brooding’, and ‘rodent run’ for distraction behaviors (Gochfeld, 1984). We classified a given species in terms of performance of distraction behavior (0 = does not perform; 1 = performs) if the paper described those behaviors as anti-predator behaviors performed during the breeding season. We obtained data for 169 species from 87 papers (Appendix). We also added data for three species (Kentish plover, *Charadrius alexandrinus*, 6 nests; little ringed plover, *C*. *dubius*, 9 nests; and little tern, *Sternula albifrons*, up to 892 nests) based on direct behavioral observations undertaken at the Morigasaki Water Recycling Center in Japan (35°57.1’S 139°75.3’W) between May and July 2017. In order to control the sampling bias by the number of papers and observations, we used the number of papers retrieved by Google Scalar as an explanatory variable. The searched word was “anti-predator”, “behavior”, and scientific name.

Data on nesting sites and colonies were obtained from del Hoyo et al. (1998). Data on body mass were obtained from Dunning (2007). We calculated the body mass of sexual dimorphic species by sex, and used the body mass data of individuals that incubate eggs (del Hoyo et al. 1998; Paton et al. 1994; Székely and Reynolds 1997; Székely et al. 2000). In cases where both sexes defend eggs, we used the average of the male and female weights. We also used the average of the male and female weights if there were no data on the sexes that defend eggs,. We obtained 100 phylogenetic trees of the study species from http://birdtree.org/ (Jetz, Thomas, Joy, Hartmann, & Mooers, 2012).

### Comparative analysis

We performed all analysis in R software (ver. 3.5.3; R Development Core Team 2019). We estimated *D*, a phylogenetic signal for discrete traits (Fritz & Purvis, 2010), for the presence of distraction behaviors using the “caper” package (Orme, 2013). We used 100 phylogenetic trees and estimated 100 *D* values for each. For all trees, we found that the phylogenetic signals of this behavior differed significantly from random (*p* < 0.001).

We used phylogenetic generalized linear mixed models with the MCMCglmm (Hadfield, 2010) and mulTree packages (Guillerme & Healy, 2014). We included the 100 phylogenetic trees as a random effect to control for the effect of phylogeny. Markov chain Monte Carlo (MCMC) simulations were run for 240,000 iterations with a 40,000 iteration burn-in period. Prior was uniform default distribution. The posterior distribution was estimated based on samples drawn after every 100 iterations.

We included three species-specific explanatory variables in the model: body mass (log-transformed; in grams), coloniality (colonial *vs*. solitary; categorical variable with two levels), nesting site (in trees or on cliffs *vs*. ground; categorical variable with two levels) and sampling bias (number of publications). We ran models in which whether a given species performed distraction was set as an independent variable.

## Results

Distraction behavior were noted in 55.62% (94/169) of the species, respectively, and evolved, and were lost multiple times, independently (Fig. 1).

**Fig. 1.**
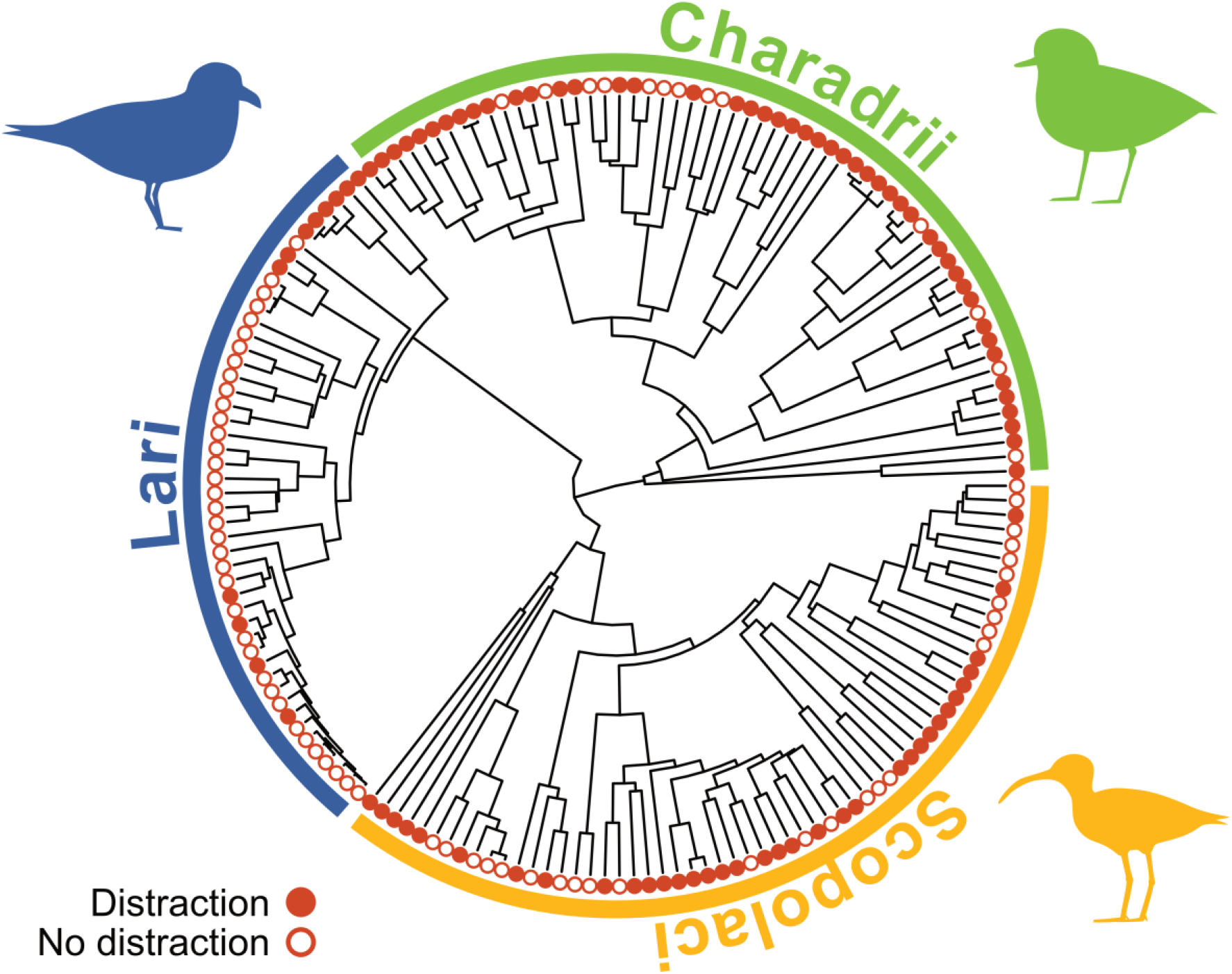
Presence of distraction behavior layered on a phylogenetic tree. Solid, performs; white, does not perform.

Species with a smaller body mass were more likely to perform distraction behavior than larger species (Table 2, Fig. 2b). In addition, solitary-nesting species commonly performed distraction behavior (78/111 species), while species that colonial-nesting did not so (18/58 species; Table 2) Although distraction behavior was observed only in one species nesting on cliffs or in trees (1/17 species), the nesting site was not statistically associated with the occurrence of distraction behavior (93/152 ground-nesting species).

**Fig. 2.**
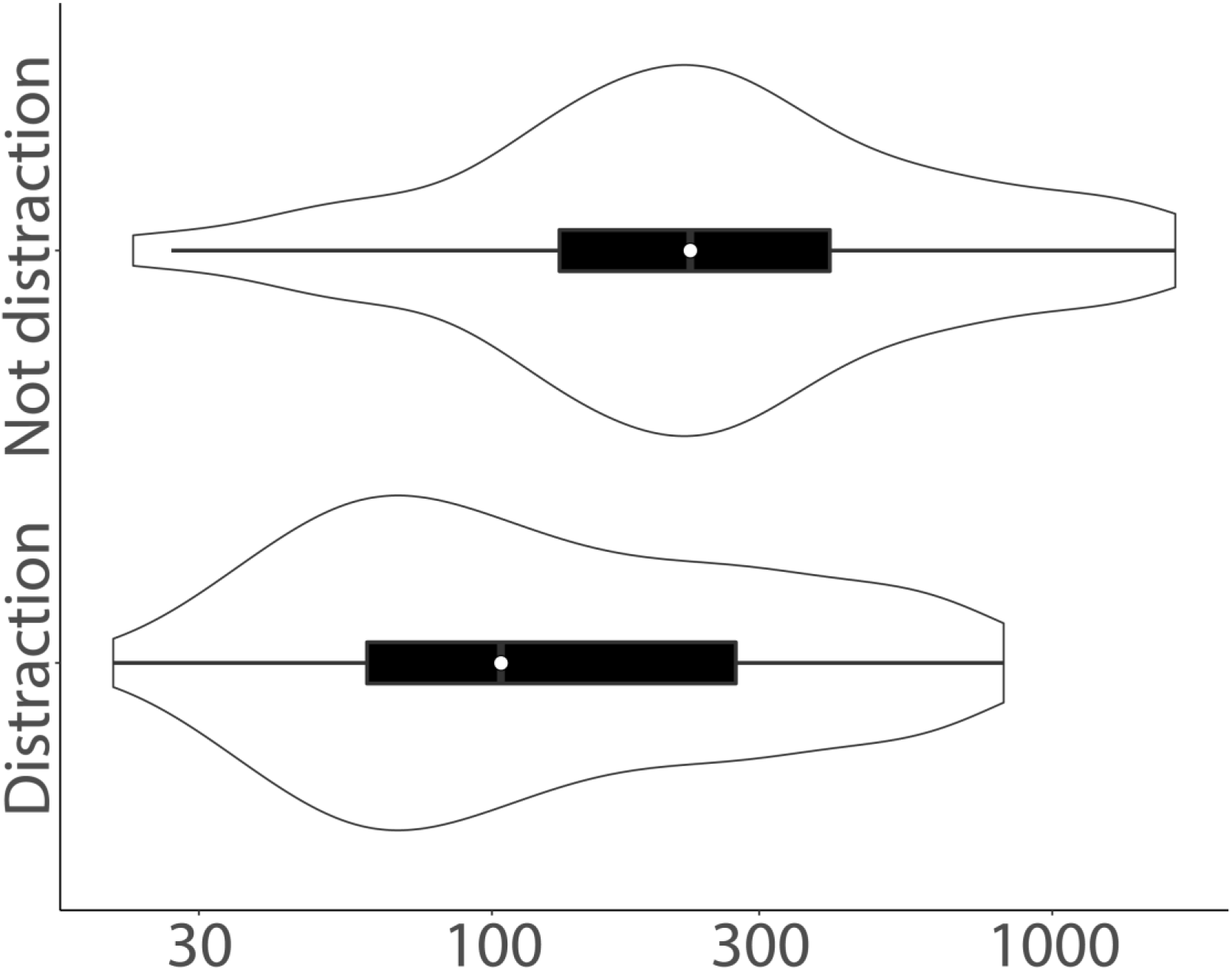
Effect of body mass (log-transformed; in grams) on distraction behavior. The number of species is indicated by the thickness of the violin plot.

## Discussion

Our analyses suggested that distraction behavior evolved, and were lost, multiple times independently, although the phylogenetic signal was significant. This suggests that species-specific patterns of anti-predator behavior are evolutionarily flexible.

Contrary to our prediction, our analysis did not show a significant effect of nest site on distraction behavior. We do not know why we failed to detect an effect of nest site, but it might be that an evolutionary transition of nest site (*i*.*e*., ground-nesting *vs*. others) was not associated with an evolutionary change in distraction; in other words, distraction may have evolved within a clade in which all species nest on the ground.

As predicted, body mass explained the inter-specific variation in distraction behavior. First, We found that small species performed distraction behavior more frequently than large species. Distraction behavior may have evolved as a substitute for attack behavior. In line with this idea, it is believed that aggressive nest defense carries a higher risk of predation for parents than distractive nest defense (D. H. Brunton, 1986; Gochfeld, 1984; Gómez-Serrano & López-López, 2017; Humphreys & Ruxton, 2020; Sordahl, 1990b).

As predicted, we found that coloniality explained the inter-specific variation in distraction behavior, with colonial species are rare to perform distraction behavior. Previously, Larsen et al. (1996) analyzed factors affecting the performance of aggressive nest defense across species, and found a relationship between aggressive defense and coloniality. Colonial species can attack a predator effectively as a group (Hoogland & Sherman, 1976; Robinson, 1985). By contrast, we found that distraction behavior was common in solitary-nesting species. Distraction behavior is as an alternative strategy because solitary species cannot do effective attacks.

In summary, this is the first study to analyze inter-specific variation in the repertoire of anti-predator behavior during the breeding season using phylogenetic comparative analyses. Although we succeeded in showing some determinants of anti-predator behavior, our analyses had several limitations. First, it is true that birds change anti-predator behavior type according to the type of predator, nesting season, nesting stage, parental sex, and condition of the parents (Andersson, Wiklund, & Rundgren, 1980; D. Brunton, 1990; Burger et al., 1989; Byrkjedal, 1987, 1989; Caro, 2005; Courchamp, Clutton-Brock, & Grenfell, 1999; Ghalambor & Martin, 2001; Gochfeld, 1984; Humphreys & Ruxton, 2020; Redondo, 1989; Sordahl, 1990a; Vincze et al., 2017). We could not control for these factors, as we focused on species-specific characteristics of the anti-predator behavior. Second, predation pressure on shorebirds varies with latitude. It is known that species nesting in high latitude area suffer lower predation risk than low latitude one (Kubelka et al., 2018; McKinnon et al., 2010). Therefore, the latitude of the nesting site may affect anti-predator behavior. This study, however, did not examined the effect of latitude because of multi-collinearity to body weight. That is, it is known that species breeding in higher latitudes have larger bodies (Olson et al., 2009). Third, our database could have underestimated the occurrence of distraction behavior, as we treated papers that did not describe anti-predator behavior as showing an absence of anti-predator behavior. The accumulation of detailed behavioral data for individual species, and comparative analyses based on an updated database, will be necessary to elucidate the details of interspecific variation in anti-predator behavior in shorebirds. Similar studies should also examine the determinants of anti-predator behavior in taxa in which multiple types of anti-predator behavior have evolved.

**Table 1.** Phylogenetically controlled MCMC generalized linear mixed models examining the effect of body mass (log-transformed; in grams), nesting site (0 = trees or cliffs, 1 = ground) and coloniality (0 = solitary, 1 = colonial) of distraction behavior. Means and 95% credible intervals (CIs; in parentheses) of the posterior distribution are shown. Results in bold are statistically significant.

|  | Posterior mean Z | 95% CI |
| --- | --- | --- |
| Intercept | 1.158 | 0.512 to 1.805 |
| Body mass | -0.120 | -0.217 to -0.021 |
| Nesting site | 0.192 | -0.051 to 0.438 |
| Coloniality | -0.218 | -0.412 to -0.024 |
| Sampling bias | 0.000 | -0.001 to 0.001 |
| Random effect: Phylogeny | 0.133 | 0.032 to 0.299 |

## Acknowledgments

We thank the members of our laboratory for useful discussions and comments on the research plan and analyses. We thank Yamashina Institute for Ornithology for permission to do literature survey.

## Compliance with ethical standards

### Ethical approval

Ethical approval is not applicable to the literature survey. Behavioral observation at the Morigasaki Water Recycling Center was conducted with the permission from the Ministry of the Environment of Japan and the Tokyo Metropolitan Government Bureau of Environment. This study was conducted under Japanese laws and the guidelines of Japanese Ethological Society.

